# Misinformation, Perceptions Towards COVID-19 and Willingness to be Vaccinated: A Population-Based Survey in Yemen

**DOI:** 10.1101/2021.02.25.432838

**Authors:** Ahmad Naoras Bitar, Mohammed Zawiah, Fahmi Y. Al-Ashwal, Mohammed Kubas, Ramzi Mukred Saeed, Rami Abduljabbar, Ammar Ali Saleh Jaber, Syed Azhar Syed Sulaiman, Amer Hayat Khan

**Affiliations:** Discipline of Clinical Pharmacy, School of Pharmaceutical Sciences, Universiti Sains Malaysia, Penang, Malaysia; Department of Pharmacy Practice, College of Clinical Pharmacy, University of Al Hodeida, Al Hodeida, Yemen; Clinical Pharmacy Department, University of Science and Technology Hospital (USTH), Sana’a, Yemen; Pharmacy Practice Department, Kulliyyah of Pharmacy, International Islamic University Malaysia (IIUM), Kuantan, Pahang, Malaysia; Department of Clinical Pharmacy and Pharmacy Practice, University of Science and Technology (UST), Sana’a, Yemen; Department of Pharmaceutical sciences, School of Pharmacy, The University of Jordan, Amman, Jordan; Department of Biopharmaceutics and Clinical Pharmacy, School of Pharmacy, The University of Jordan, Amman, Jordan; Dept of Clinical Pharmacy & Pharmacotherapeutics, Dubai Pharmacy College for Girls, Dubai, UAE; Advanced Medical and Dental Institute, Universiti Sains Malaysia, Kepala Batas, Penang, Malaysia

**Keywords:** COVID-19, Misinformation, Vaccine acceptance, Perception, Severity, Susceptibility

## Abstract

**Background:** Since the beginning of the COVID-19 outbreak, many pharmaceutical companies were racing to develop a safe and effective COVID-19 vaccine. Simultaneously, rumors and misinformation about COVID-19 were and still widely spreading. Therefore, this study aimed to investigate the prevalence of COVID-19 misinformation among the Yemeni population and its association with vaccine acceptance and perceptions.

**Methods:** A cross-sectional online survey was conducted in four major cities in Yemen. The constructed questionnaire consisted of four main sections (sociodemographic data, misinformation, perceptions (perceived susceptibility, severity and worry), and vaccination acceptance evaluation). Subject recruitment and data collection were conducted online utilizing social websites and using the snowball sampling technique. Descriptive and inferential analyses were performed using SPSS version 27.

**Results:** The total number of respondents was 484. Over 60% of them were male and had a university education, more than half had less than 100$ monthly income and were Khat chewers, while only 18% were smokers. Misinformation prevalence ranged from 8.9% to 38.9%, depending on the statement being asked. Men, university education, higher income, employment, and living in urban areas were associated with a lower misinformation level (*p* <0.05). Statistically significant association (*p* <0.05) between university education, living in urban areas, and being employed with perceived susceptibility were observed. The acceptance rate was 61.2% for free vaccines, but it decreased to 43% if they had to purchase it. Females, respondents with lower monthly income, and those who believed that pharmaceutical companies made the virus for financial gains were more likely to reject the vaccination (*p* <0.05).

**Conclusion:** The study revealed that the acceptance rate to take a vaccine was suboptimal and significantly affected by gender, misinformation, cost, and income. Furthermore, being female, Non-university educated, low-income, and living in rural areas were associated with higher susceptibility to misinformation about COVID-19. These findings show a clear link between misinformation susceptibility and willingness to vaccinate. Focused awareness campaigns to decrease misinformation and emphasize the vaccination’s safety and efficacy might be fundamental before initiating any mass vaccination in Yemen.

## Introduction

More than one year has passed since the begging of the severe acute respiratory syndrome coronavirus-2 (SARS-CoV-2) pandemic. The first report of COVID-19 by the World health organization (WHO) was made on the 31^st^ of December 2019, and by the 11^th^ of March, 2020, a global pandemic was declared [1]. In a systematic review, out of 53000 hospitalized COVID-19 patients, more than 20% suffered from severe and dangerous symptoms, and the mortality rate was 3.1% [2]. Protective measures such as wearing a facial mask, avoiding crowded areas, limiting direct contact, and maintaining a safe distance proved to be effective in restraining the spread of the disease [3,4]; however, they do not present a long term solution of COVID-19 pandemic and the development, and the usage of an effective vaccine will be essential for controlling the current pandemic as epidemiologists projected the pandemic to last beyond 2021 based on mathematical modeling [5]. More than 30 biotech and pharmaceutical companies were racing to develop a safe COVID-19 vaccine; at least three vaccines were approved worldwide, AstraZeneca, Pfizer-Biotech, and Moderna [6]. Large-scale vaccination programs are planned to reach herd immunity against COVID-19; however, such a program’s success will majorly depend on the public response toward the vaccine.

There are vital factors that might influence the public response and acceptance of the newly developed vaccines, like perceived susceptibility and severity towards COVID-19 and the misinformation spread. To correct misinformation, we should first identify what the public believes and what is right and wrong. Understanding the nature of misinformation and exploring the source of inaccurate information can promote behavior change interventions developed to solve the problem and provide accurate information [7]. Misinformation can lead to serious consequences such as increased burden on the health care system due to emergency hospitalization caused by false beliefs or wide speared of wrong health advice, and the spread of politically motivated conspiracies like 5G masts can cause COVID-19, or the vaccine is designed to control the humanity or reduced the population; due to that, some people burnt 5G masts while others are reluctant to take the vaccine [8]. Furthermore, a recent study has shown that a viral YouTube video about COVID-19, with more than 62 million views, contained 25% of misleading information [9]. Some evidence also indicates that misconceptions are more common than anticipated, with 46% of the British being exposed to fake news about coronavirus [8].

Doubts about the vaccine effectiveness, safety, and usefulness are also major obstacles for the population acceptance of COVID-19 vaccines worldwide. For example, in France, 25% of the population reported refusing the vaccine due to safety concerns [10]. In Saudi Arabia, 36% showed no interest in the vaccine [11], while in the USA, the FDA’s emergency use authorization was associated with a lower probability of accepting the vaccine [12]. The willingness to be vaccinated was linked to more positive general vaccination beliefs and attitudes and weaker beliefs about the vaccine being unsafe or it causes severe side effects. Also, well-informed subjects had a positive perception about the benefits of the vaccine and the danger of COVID-19; furthermore, subjects previously vaccinated against influenza had a more positive perception of the vaccine [13]. Vaccination acceptance can reflect the general perception, attitude, and beliefs about the newly developed vaccine, and the increased awareness of the risk of COVID-19 and the benefits of vaccination can increase the general uptake of the vaccine.

To the best of our knowledge, no previous research in the middle east assessed the association between misinformation and willingness to vaccinate. In addition, the conducted research about COVID-19 in Yemen is exceptionally scarce. In a recent study, we have found that the lack of awareness about COVID-19 among the public was the most common obstacle that undermines all efforts to control the outbreak [14]. Accordingly, in this study, we tried to investigate the prevalence of COVID-19 misinformation among the Yemeni general population and its association with vaccine acceptance and perception. Also, factors affecting the perceptions, misinformation, and willingness to vaccinate were evaluated.

## Methods

### Study design and settings

This study is a cross-sectional online survey in Yemen. Subject recruitment and data collection were conducted online using the snowball sampling technique. Data were collected in the first two weeks of the outbreak in Yemen from April 12^th^, 2020, until the April 26^th^, from four major cities in Yemen: Sana’a, Al-Hudaidah, Ta’aiz, and Aden. In each city, the response was from both urban and rural households. All included subjects were above 18 years. Social media platforms like WhatsApp and Facebook were used to distribute google form questionnaires. Subjects were recruited in the study using the simplified snowball sampling technique, and they were requested to pass the invitation to their contacts; the estimated time to complete the survey was around 10 min.

### Ethical approval

This study was part of a project about COVID-19 in Yemen. The project was registered with the ethical committee of the Medical Research, University of Science and Technology, Sana’a, Yemen, with the following number: ECA/UST189. An electronic consent statement to be ticked by all participants who agreed to participate, those who did not tick it will not be able to fill the questionnaire.

### Data Collection Tool

The constructed questionnaire (S1 file) was divided into four sections; Section A: demographics data such as age, gender, marital status, residential area, medical insurance, education level, work nature, presence of chronic disease, smoking, and khat chewing status. Section B: Misinformation about COVID-19. This section contains seven statements with five possible answers on a 5-Likert scale (Strongly disagree to agree strongly) about commonly spread misinformation at the time of data collection. The scores for each statement ranged from 1 to 5, and the overall from 7 to 35, a higher score, indicated a higher level of misinformation. For inferential analysis, misinformation was categorized using the median as misinformed (>19) and informed (≤19). Section C: COVID-19 perception, which was divided into three subsections; perceived susceptibility, perceived severity/threat, and perceived worry. Perceived susceptibility is the individual’s belief of the chances to get a COVID-19. It contained four items with four possible choices (not at all likely, slightly likely, somewhat likely, and very likely), and the total score ranged from 4 to 16. For perceived severity, belief about how serious or dangerous the COVID-19 and its consequences are, a 4-Likert scale ranged from not dangerous at all to very dangerous was used, and the total score ranged from 5 to 20. The perceived worry (5 items) was assessed on a 4-Likert scale (not at all worried, slightly worried, moderately worried, and very worried), and the total score ranged from 5 to 20. The final score for perception subscales was categorized into perceived and not perceived using the median for Chi-Square analysis.

Section D: two questions, with yes, no, not sure answers, focused on the vaccine acceptance among Yemeni people in two situations, a free effective vaccine and a vaccine with a cost of about 10 thousand Yemeni Rials (approximately 15$ at the time of study period). The constructed questionnaire was structurally designed to be based on self-reporting, and some of its’ components were adapted from previously published studies about the Ebola virus [15,16].

The designed questionnaire’s contents were checked, evaluated, and validated by a panel of experts with academic, clinical, and questionnaire construction backgrounds. The questionnaire was checked for reliability using Cronbach’s Alpha. The results from Cronbach’s Alpha were as following; COVID-19 misinformation (0.647), perceived susceptibility (0.870), perceived severity (0.757), and perceived worries (0.830).

### Sample Size

A sample size of 385 was estimated using the Daniel formula [17] with an expected prevalence of 50% for misinformation and vaccination acceptance, to have the highest number of respondents [18], using a confidence interval of 95% and precision of 0.05 and estimated population of 30 million in Yemen [19]. We recruited additional 100 subjects to reach a final sample of 484 and get at least 80% power.

### Statistical Analysis

Descriptive statistics were used to analyze the data’s sociodemographic characteristics and the responses to questions concerning misinformation, perceptions, and confidence acceptance of the COVID-19’s vaccines. Frequencies and percentages were used to present categorical variables. A Pearson Chi-Square was conducted to detect the association between respondents’ characteristics and their misinformation, perception, and vaccination acceptance. Statistical analysis was done using the Statistical Package for the Social Sciences (SPSS) (Version 27.0; IBM corp). P<0.05 was taken as a cut point for statistically significant results.

## Results

The total number of subjects who participated in this study was 484. Approximately 61% of whom were male and had a university education. While more than half of the participants were Khat chewers, only 18% were smokers. Also, the majority of participants (85%) were free from chronic diseases. In addition, more than half of the participants had less than 100$ income, and one-third of respondents were unemployed (Table 1).

**Table 1.**
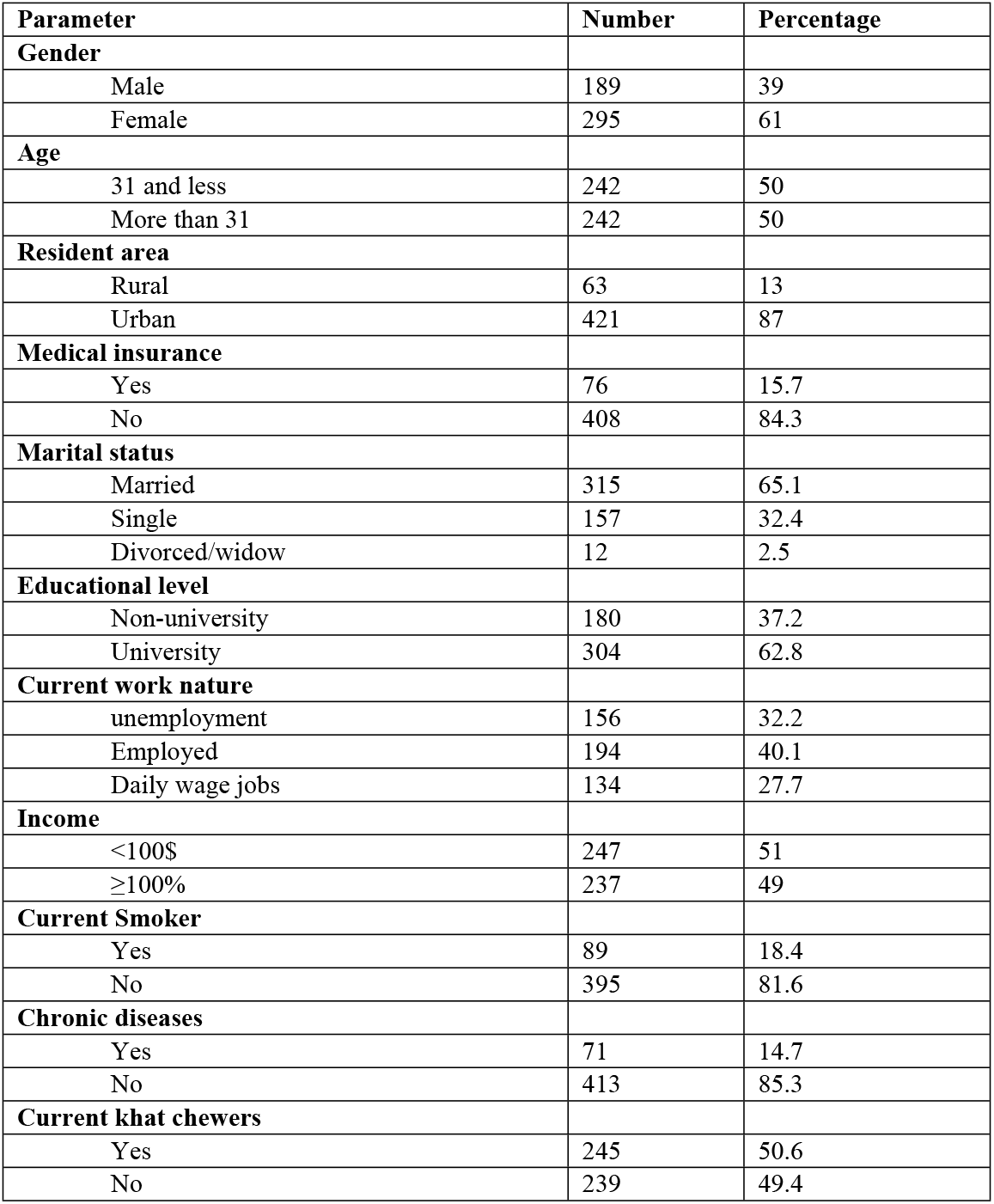
Socio-demographic characteristics.

### Misinformation

For the prevalence of misinformation, slightly less than a quarter of participants (23.2%) agree or strongly agree that COVID-19 was a human-made virus designed by pharmaceutical companies for financial gains, while almost two fifths (40%) saw the virus as a human-made biological weapon (Table 2). The proportion of respondents who agreed or strongly agreed that COVID-19 could not be transmitted in hot weather stood just under a third (30%), and almost the same proportion thought that most people who are infected with coronavirus would die. Regarding preventing the viral infection, around 30% of subjects thought that COVID-19 infection could be prevented or treated using natural remedies like hot anise tea or eating raw garlic. Notably, a small minority of respondents (9%) had the misconception that antibiotics could treat COVID-19 infections. Upon assessing the association between sociodemographic factors and misinformation, we found that men, university education, higher income, being employed, and living in urban areas are significantly associated (*p* <0.05) with being informed (lower level of misinformation) (Table 4).

**Table 2.** Covid-19 misinformation.

| Item | Strongly disagree | Disagree | Neutral | Agree | Strongly agree |
| --- | --- | --- | --- | --- | --- |
| COVID-19 is human-made for pharmaceutical companies' financial gains | 68 (14) | 100 (20.7) | 204 (42.1) | 71 (14.7) | 41 (8.5) |
| COVID-19 was created by human as a biological weapon | 43 (8.9) | 64 (13.2) | 189 (39) | 134 (27.7) | 54 (11.2) |
| COVID-19 virus cannot be transmitted in areas with hot climates | 41 (8.5) | 125 (25.8) | 152 (31.4) | 154 (31.8) | 12 (2.5) |
| Children will not be infected or carry the virus | 105 (21.7) | 207 (42.8) | 98 (20.2) | 68 (14.0) | 6 (1.2) |
| Most people who get the coronavirus will die | 90 (18.6) | 179 (37.0) | 61 (12.6) | 126 (26.0) | 28 (5.8) |
| COVID-19 can be prevented or treated by eating raw garlic and drinking hot tea containing anise | 49 (10.1) | 129 (26.7) | 160 (33.1) | 132 (27.3) | 14 (2.9) |
| Antibiotics are effective in preventing and treating the new coronavirus | 145 (30.0) | 147 (30.4) | 149 (30.8) | 41 (8.5) | 2 (0.4) |

### Willingness to vaccinate

Around two-fifths of respondents (61 %) would take the vaccine if they were offered a free one (Fig. 1.A). However, the vaccination acceptance decreased to 43% if they had to purchase it (Fig. 1.B). Regarding the reasons that make the participants reluctant to be vaccinated. Around a third of participants (31.1%) were concerned about the vaccine safety and side effects, 21.5% were skeptical of the vaccine’s efficacy, 24.1% were concerned about the vaccine’s price, while only 5% stated that they did not need it as they had no risk for COVID-19 (Fig 1.C). For factors affecting the acceptance of the vaccine, men were more likely to accept the vaccine than women, whether provided for free or to purchase it (Free: 66.4% vs 52.9%; respectively, *p* < 0.05, purchase: 47.5% vs 36%; respectively, *p* < 0.05) (Table 4). Also, respondents who were above 31 years of age were more willing to be vaccinated for free than the younger group (69.4% vs 52.9%; respectively, *p* <0.001), while the data showed no significant difference for the purchased vaccine. Moreover, employed individuals were more willing to purchase a vaccine than unemployed and daily wages workers (49.5%, 36.5%, vs 41%; respectively, *p* < 0.05). Furthermore, participants who chewed Khat had a higher acceptance for the vaccine regardless of cost status (Table 4). Notably, respondents with lower monthly income (less than 100$) were more likely to reject the vaccination whether provided for free (48.6% vs 28.7%; respectively, *p* < 0.001) or with a cost of 15$ (70% vs 43.5%; respectively, *p* < 0.001).

**(Fig 1. A-C).**
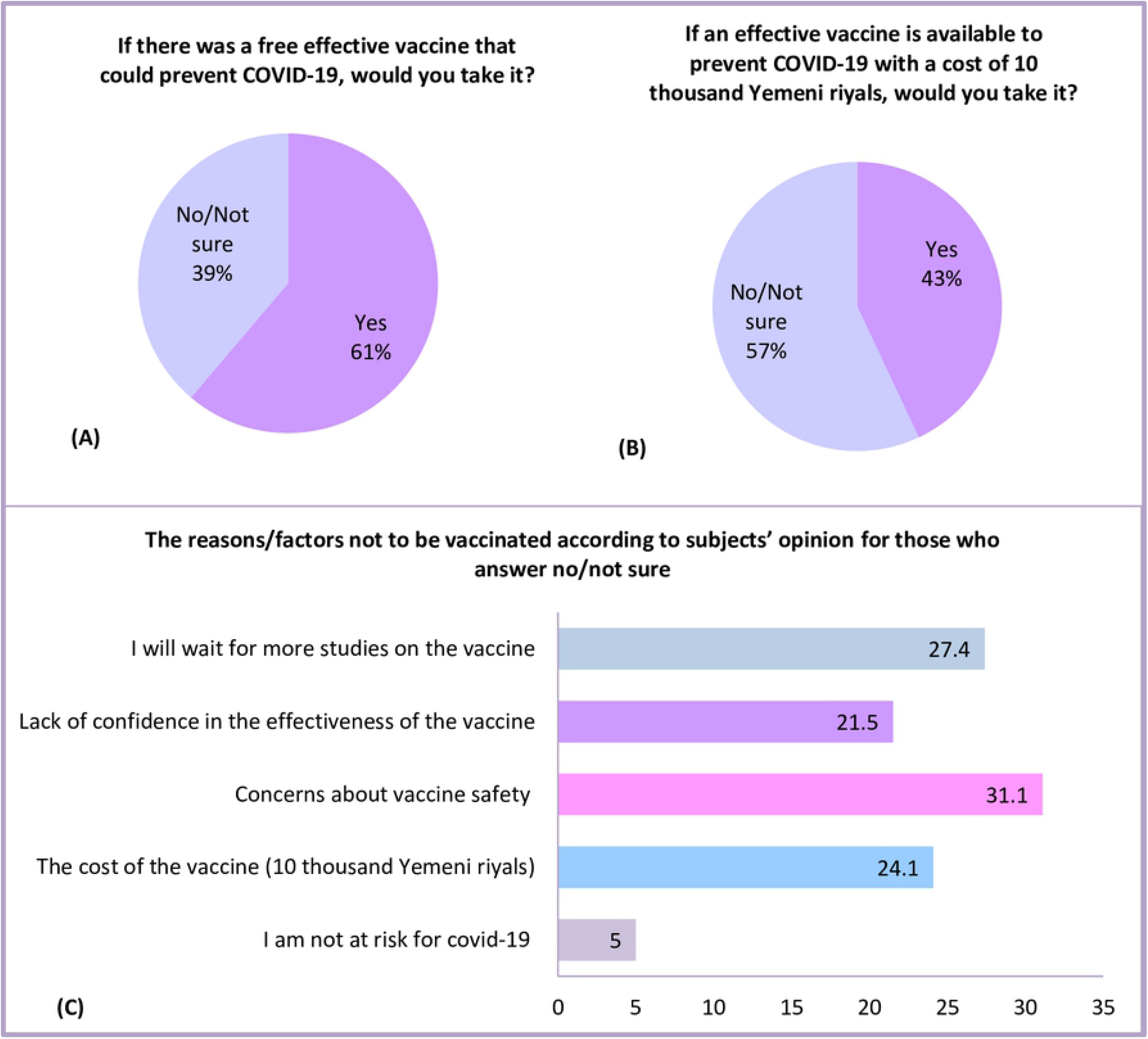
Willingness to vaccinate and barriers hindering Yemeni people from vaccination

### Perceived susceptibility

Only 20.9% of respondents believed it was somewhat likely or very likely that they would be infected with COVID-19 (Table 3). Also, less than one-fifth of participants believed it was somewhat likely or very likely that their family would get COVID-19 infection. This indicates a low perception of self and family susceptibility among the majority of the respondents. Interestingly, when asked about their city and governorate, the perceived susceptibility increased to 35.4% and 42.6%, respectively. Statistically significant association (*p* <0.05) between university education, living in the urban areas, and being employed with perceived susceptibility was observed in table 4.

**Table 3.** Perception of the community towards COVID-19.

| **Perceived susceptibility** | **Not at All Likely** | **Slightly Likely** | **Somewhat Likely** | **Very Likely** |
| --- | --- | --- | --- | --- |
| How likely is it that you will be infected with COVID-19 within the next few months? | 178 (36.8) | 205 (42.4) | 76 (15.7) | 25 (5.2) |
| How likely is it that one of your family will be infected with COVID-19 within the next few months? | 177 (36.6) | 216 (44.6) | 68 (14.0) | 23 (4.8) |
| How likely is the onset of the COVID-19 outbreak in your city within the next few months? | 95 (19.6) | 218 (45.0) | 87 (18.0) | 84 (17.4) |
| How likely is the onset of the COVID-19 outbreak in your governorate within the next few months? | 65 (13.4) | 213 (44.0) | 100 (20.7) | 106 (21.9) |
| **Perceived severity/threat** | **Not dangerous at all** | **Slightly dangerous** | **Moderately dangerous** | **Very dangerous** |
| In general, how dangerous do you think the COVID-19 pandemic is? | 5 (1.0) | 34 (7.0) | 108 (22.3) | 337 (69.6) |
| How dangerous do you think it would be if you are diagnosed with COVID-19? | 7 (1.4) | 47 (9.7) | 110 (22.7) | 320 (66.1) |
| How dangerous do you think would COVID-19 be on you if it begins to spread to your community? | 32 (6.6) | 99 (20.5) | 156 (32.2) | 197 (40.7) |
| How dangerous do you think it would be on your city if COVID-19 starts spreading in your governorate? | 10 (2.1) | 21 (4.3) | 66 (13.6) | 387 (80.0) |
| How dangerous do you think the consequences of COVID-19 disease would be on your country? | 8 (1.7) | 13 (2.7) | 58 (12.0) | 405 (83.7) |
| **Perceived Worry** | **Not at all worried** | **Slightly worried** | **Moderately worried** | **Very worried** |
| How worried are you about COVID-19 at this moment? | 69 (14.3) | 211 (43.6) | 93 (19.2) | 111 (22.9) |
| How worried are you that you will be infected with COVID-19 in the next few months? | 96 (19.8) | 223 (46.1) | 72 (14.9) | 93 (19.2) |
| How worried are you that someone you know (relatives, friends, etc) will be infected with COVID-19 in the next few months? | 39 (8.1) | 166 (33.3) | 99 (20.5) | 185 (38.2) |
| How worried are you that an outbreak of COVID-19 will happen in your city? | 13 (2.7) | 99 (20.5) | 79 (16.3) | 293 (60.5) |
| How worried are you that you will not be able to go outside of your house during an outbreak happen in your city? | 73 (15.1) | 121 (25.0) | 84 (17.4) | 206 (42.6) |

**Table 4.** Association between demographics and subjects’ perception, vaccine acceptance, and misinformation.

| **Parameter** | **Susceptibility(Perceived, N=223)** | **Severity(Perceived, N=234)** | **Worry(Perceived, N=216)** | **Free Vaccine (Accepted, N=296)** | **Purchased vaccine (Welling, N=208)** | **Misinformation (Informed, N=253)** |
| --- | --- | --- | --- | --- | --- | --- |
| **Gender** |  |  |  |  |  |  |
| Male | 142(51.9) | 137(51.3) | 129(46) | 196(66.4)* | 140(47.5)* | 168(56.9)* |
| Female | 81(57.1) | 97(46.4) | 87(43.7) | 100(52.9) | 68(36) | 85(45) |
| **Age** |  |  |  |  |  |  |
| Less than 31 | 121(54.3) | 128(43.8) | 98(48.8) | 168(52.9) | 109(40.9) | 134(49.2) |
| More than 31 | 102(45.7) | 106(52.9)* | 118(40.5) | 128(69.4)*** | 99(45) | 119(55.4) |
| **Education** |  |  |  |  |  |  |
| University | 159(52.3)*** | 135(57.7)* | 125(50.6)* | 193(63.2) | 125(41.1) | 178(61.5)*** |
| Non-university | 64(35.6) | 99(42.3) | 91(41.1) | 104(57.8) | 83(46.1) | 66(36.7) |
| **Residential Area** |  |  |  |  |  |  |
| Urban | 205(48.7)* | 189(44.9)*** | 186(44.2) | 254(60.3) | 176(41.8) | 229(54.4)* |
| Rural | 18(28.6) | 45(71.4) | 30(47.6) | 42(66.7) | 32(50.8) | 24(38.1) |
| **Medical Insurance** |  |  |  |  |  |  |
| Yes | 41(53.9) | 30(39.5) | 33(43.4) | 47(61.8) | 38(50) | 45(59.2) |
| No | 182(44.6) | 204(50) | 183(49.9) | 249(61) | 170(41.7) | 208(51) |
| **Work** |  |  |  |  |  |  |
| Employed | 104(53.6)* | 85(43.8) | 83(42.8) | 131(67.5) | 96(49.5)* | 124(63.9)*** |
| Unemployed | 73(46.8) | 73(46.8) | 73(46.8) | 90(57.7) | 57(36.5) | 74(47.4) |
| Daily Wages | 46(34.3) | 76(56.7) | 60(44.8) | 75(52) | 55(41) | 55(41) |
| **Income** |  |  |  |  |  |  |
| <100\$ | 109(44.1) | 131(53) * | 117(47.4) | 127(51.4) | 74(30) | 94(38.1) |
| ≥100\$ | 114(48.1) | 103(43.5) | 99(41.8) | 169(71.3) *** | 134(56.5) *** | 159(67.1) *** |
| **Smoking** |  |  |  |  |  |  |
| Yes | 45(50.6) | 45(50.6) | 37(41.6) | 55(61.8) | 42(47.2) | 40(54.9) |
| No | 178(45.1) | 191(47.8) | 179(45.3) | 241(61) | 166(42) | 213(53.9) |
| **Khat Chewing** |  |  |  |  |  |  |
| Yes | 117(47.8) | 126(51.4) | 109(43.7) | 162(66.1)* | 116(47.3)* | 127(51.8) |
| No | 106(44.4) | 108(45.2) | 105(45.6) | 134(56.1) | 92(38.5) | 126(52.7) |
| **Chronic Conditions** |  |  |  |  |  |  |
| Yes | 31(43.7) | 47(66.2)** | 42(59.2)** | 49(69) | 32(45.5) | 33(46.5) |
| No | 192(46.5) | 184(45.3) | 174(42.1) | 247(59.8) | 176(42.6) | 220(53.3) |

Chi-Square test was used to examine the association. \* p-value <0.05, \*\* p-value <0.01, \*\*\* p-value <0.001.

### Perceived severity/threats

The perceived threat data indicated that around 70% of respondents thought of COVID-19 as a dangerous disease. However, only two-fifths perceived COVID-19 as very dangerous to them if it started spreading in their community. Remarkably, the perceived severity increased among the respondents when they were asked about how dangerous COVID-19 would be on their city, and its consequences would be on the country, with 80% and 83% saw it as very dangerous, respectively. Respondents who were above 31 years of age, had low income (<100$), and those who were living in the urban area or with a chronic disease condition are more likely to have a perception of the disease’s severity (Table 4).

### Perceived worry

More than half of the respondents (57.9%) were not worried at all or slightly worried about the COVID-19 situation. Moreover, just over a third of subjects (34.1 %) were at least moderately worried about attracting the infection themselves in the next few months. This percentage of perceived worry increased to more than half (58.7%) when respondents were asked about their relatives and friends. While the majority of people (76.8%) were moderately worried or very worried that an outbreak would happen in their cities, less percentage of them (58%) were worried about the restrictions that might come with a sudden outbreak, like being unable to go out. Perceived worry was significantly associated (*p* <0.05) with respondents who had a university education or had a chronic disease condition (Table 4).

### The effect of misinformation on perceptions and willingness to vaccinate

The data showed that respondents who believed that pharmaceutical companies made the virus for financial gains had a significantly lower acceptance rate for a free vaccine (50.9% vs 64.2%; respectively, *p* < 0.05) or with a cost of 15$ (34.8% vs 45.5%; respectively, *p* < 0.05) compared to those who did not think so (Table 5). The participants who believed that most people with COVID-19 would die had significantly higher perception for severity (60.4% vs 42.7%; respectively, *p* < 0.001) and worry for the disease (57.1% vs 38.8%; respectively, *p* < 0.001). Interestingly, people who thought that COVID-19 could be prevented by eating garlic had a significantly lower level of susceptibility perception for COVID-19 (39% vs 49.1%; respectively, *p* < 0.05). The rest of the misinformation statements showed no significant association with perceived susceptibility, severity, worry, and vaccine acceptance.

**Table 5.**
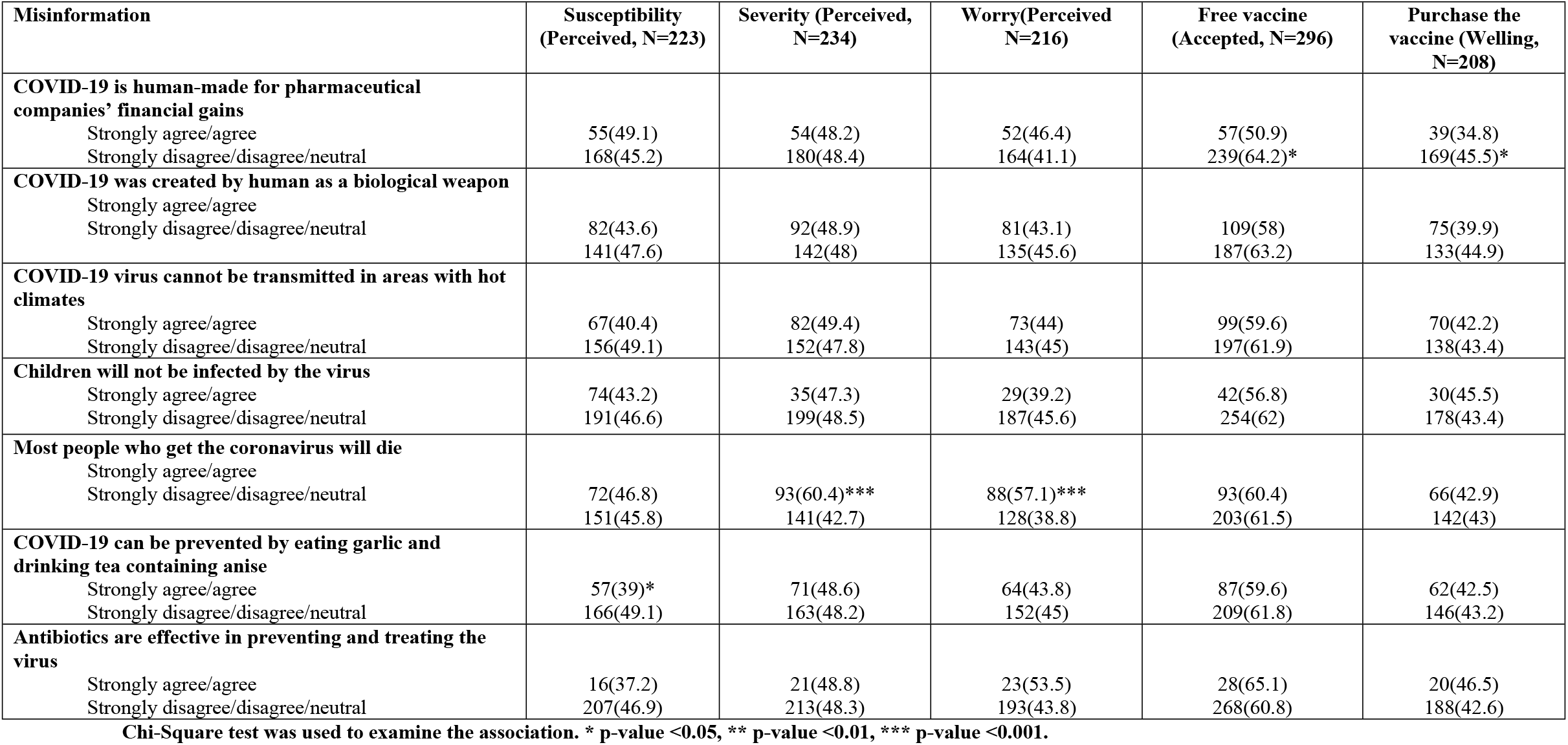
Association between subjects’ misinformation and the perception and the vaccine acceptance of the vaccine.

## Discussion

Overall, misinformation was not high among Yemenis except for the misconception that humans have created COVID-19 as a biological weapon, where approximately two-fifths believed so. This was consistent with a recent finding from Nigeria, where 39% believed that COVID-19 was part of biological warfare [20]. A higher percentage (57%) of the population was reported to have the same belief in Jordan [21]. Even though experts refuted the idea that COVID-19 was engineered in the laboratories, these findings suggest that this misconception is still common.

Female gender, Non-university education, low-income, and rural areas were significantly associated with being misinformed about COVID-19. These results align with those reported in Jordan, where beliefs that COVID-19 is part of a conspiracy theory and biological warfare were more common in females and among people with low education level and income [21]. Such findings could be attributed to multiple factors; for example, in Yemen, males have better access to education than females, reports from UNESCO have shown that in Yemen, the literacy among male subjects was more than 30% higher than the female, with around 85% of male subjects reaching high school [22]. Also, Rampersad and Althyabi found that gender can weakly and indirectly influence misinformation acceptance, while education showed a strong negative effect on accepting rumors and misinformation [23]. Well-educated people are less likely to accept misinformation. They depend on reliable sources and tend to search for expert opinions rather than accept misinformation, and usually, they have more analytical thinking [24]. The lower level of misinformation among urban area residents could be partially explained by the availability of better access to education and health facilities than those in rural areas due to higher population density, closer proximity, and transportation availability [25].

We found that 61 % of the public in Yemen agreed to be vaccinated against COVID-19. This finding was lower than that reported from an international survey of 19 countries which found 71.5% of participants were willing to take the vaccine [26]. The main barriers that make our participants reluctant to be vaccinated were concerns over vaccine safety, efficacy, and price. The price, which is 15 USD only, decreased the acceptance rate by 20%, suggesting that cost is a significant barrier to vaccination in Yemen, and a higher rejection rate would be expected if people in Yemen had to purchase a higher-cost vaccine. This is because Yemen is one of the middle east’s poorest countries, where almost 50% of the population lives below the poverty line, and around 20% earn only 1.2 USD per day [27,28]. Also, the association between low financial status and vaccination rejection was apparent in this study; those who had a monthly income of less than 100 USD and unemployed individuals were less likely to purchase a vaccine. Notably, the acceptance rate for a free COVID-19 vaccine was also suboptimal (61 %) and below the percentage needed to meet the anticipated levels of herd immunity, suggesting that it is vital for healthcare authorities and international organization working in Yemen not only to ensure that these populations have a free vaccine, but also that trust in the vaccine efficacy and safety is built up prior to roll-out vaccination campaigns. More efforts should be targeted to female, younger, and low-income groups as these were more likely to reject a free vaccination, in alignment with recent studies from the UK [13,29,30].

Another clear finding from this study that misinformation could influence people’s decision to get vaccinated. In this light, people who believed that COVID-19 is human-made for pharmaceutical companies’ financial gains were more likely to reject the vaccine. Such finding is concordant to other emerging studies in the context of COVID-19 that have linked specific theories of conspiracy to lower willingness to adopt behaviors in public health. Brennen et al. found that much of the misleading information about COVID-19, like the conspiracy theory is originated from fake news, which can be associated with photoshopped pictures usually used as fake pieces of evidence and can be spread easily through social media [31]. This kind of conspiracy ideas can present a significant obstacle for mass vaccination programs, and it can play a role in facilitating the virus spread because those who believe in such theories tend not to take preventive measures like social distancing and do not follow standard operating procedures [32,33]. Therefore, it is important to debunk this kind of theory and try to confront the public’s misinformation to control the virus spread and increase vaccine acceptance among the Yemeni population.

For perceptions, the participants showed a low-self and family perception regarding the susceptibility of being infected. Similarly, during the H5N1 virus outbreak in Hong Kong, it was found that the vast majority of subjects had less perception toward the risk of being infected themselves compared to the risk of a general outbreak in their cities [34]. Furthermore, people were too concerned and worried about their cities and communities in case of a major outbreak, which is harmonious with our findings. Factors such as age, employment, urban areas, and education were also associated with a better perception among the included subjects. In the United States, they have found that the perception toward the risks of COVID-19 was higher among older subjects, while the younger subjects were more worried about the pandemic; furthermore, women’s perception of risks slightly higher than men, and they were more worried than men; however, the interactions between gender and age were insignificant [35]. Interestingly, our participants’ level of perception increased (perceived susceptibility, severity, and worry) when asked about themselves, their relatives, and cities, respectively, indicating they had a higher perceived susceptibility and worry for their relatives and communities. This suggests that altruistic messaging to protect their families, friends, country might be a useful strategy among the Yemenis to increase their acceptance rate for the COVID-19 vaccine [33].

### Strength and limitations of the study

There are a few limitations to the conducted study. Responses were received mainly from four major cities in Yemen, limiting the generalization of the whole country’s findings. An online snowballing sampling technique has been used, which can impact the neutrality of subjects’ selection and recruitment in the study. Furthermore, in cross-sectional studies, the results represent only a point when the data has been collected; thus, the public’s misinformation, perceptions, vaccination acceptance can be changed over time.

Despite the presented limitations, the sample size was adequate, and the participants were from the largest cities in Yemen, which gives a close enough image and a realistic idea about the presented topic in Yemen at the first two weeks of the outbreak in Yemen. The conducted study provides a good insight into the misinformation, perceptions, and acceptance of the vaccine among Yemenis, opening the door for more comparative research and investigations to be conducted in the future or after awareness campaigns and educational interventions. The study also provides a good insight into COVID-19 vaccine acceptance in a low-income, less-developed country like Yemen. Importantly, the study findings provide useful insight for policymakers, healthcare planners, and international organizations planning to support or donate vaccines to Yemen.

## Conclusion

The study revealed that the acceptance rate to take a vaccine was suboptimal and significantly affected by gender, misinformation, cost, and income. Furthermore, being female, Non-university educated, with low-income, and living in rural areas were associated with higher susciptability to misinformation about COVID-19. Taken together, these findings show a clear link between misinformation susceptibility and willingness to vaccinate. Focused awareness campaigns to decrease misinformation and emphasize the vaccination’s safety and efficacy might be fundamental before initiating any vaccination program in Yemen.

## Supporting information

This is the S1 file title.

S1 file. Study Questionnaire

